# Shift from acquisitive to conservative plant strategies with increasing drought and temperature extremes in an alpine shrub

**DOI:** 10.1101/2025.06.26.661801

**Authors:** Dinesh Thakur, Nikita Rathore, Veronika Jandova, Zuzana Münzbergová, Jiri Dolezal

## Abstract

**Background and Aims:** Trait-based approaches have advanced our understanding of plant strategies, yet they often focus on leaf-level traits, overlooking the functional roles of stem anatomy and twig characteristics. We investigated intraspecific trait variation in *Salix flabellaris*, an alpine dwarf shrub, along climatic gradients in the Himalayas. Our goal was to identify distinct axes of trait variation related to stem, twig, and leaf traits, assess their environmental drivers, and evaluate population-specific growth responses to recent climate change.

**Methods:** We measured anatomical and morphological traits in stem, twig, and leaves across central and marginal populations along three Himalayan transects. Environmental gradients included variation in growing season temperature and soil moisture. Basal area increment from 2000 to 2021 was analyzed to assess long-term growth trends in different areas.

**Results:** Trait dimensions were largely independent, reflecting distinct ecological strategies: (1) stem anatomical trade-off between hydraulic safety and conductivity; (2) twig dimension balancing construction costs and mechanical strength; and (3) leaf dimension along the exploitative–conservative axis. Higher temperatures enhanced performance manifested as larger twigs and reduced tissue construction costs but only under sufficient soil moisture conditions. Central populations at mid-elevations displayed the favorable trait combinations and highest growth rates. In contrast, marginal populations (higher and lower elevations) showed traits indicating structural reinforcement and conservative resource use. Climate warming over recent decades enhanced stem growth primarily in high-elevation populations, where low-temperature constraints were relaxed.

**Conclusions:** This study demonstrates that stem, twig, and leaf traits represent distinct yet complementary strategies, with environmental filtering shaping their expression along climate gradients. Central populations exhibit highest growth under current conditions, while climate change is shifting growth advantages toward higher elevations. These findings highlight the need for integrated, multi-organ trait assessments to predict species performance, persistence, and potential range shifts under future climatic scenarios.

## INTRODUCTION

Climate change is profoundly impacting all levels of biodiversity, from individual organisms to entire ecosystems (Malhi et al., 2020; Thakur et al., 2023). High-elevation alpine ecosystems are particularly vulnerable as they are changing at a faster rate than many other environments (Pepin et al., 2015; Tito et al., 2020). This accelerated change poses a significant threat to the plant species that thrive in these unique settings, as well as to the overall stability, biodiversity, and functionality of these ecosystems. Thus, investigating how environmental factors influence plant functioning is essential for understanding and predicting the ecological consequences of climate change on these fragile ecosystems.

Plant functional traits are crucial for understanding plant adaptive strategies (such as conservative to acquisitive plant strategy), the effects of various factors on plants, and the ecological roles of plants within ecosystems (Kühn et al., 2021; McGill et al., 2006; Perez-Harguindeguy et al., 2013; Wright et al., 2004). Traits related to the plant resource economics spectrum, such as specific leaf area (SLA), leaf dry matter content (LDMC), specific twig length (STL), twig increment, twig density (TDen), twig volume increment (TVol), affect primary productivity, litter decomposability, soil carbon storage, and nutrient cycling (Perez-Harguindeguy et al., 2013; Smith et al., 2017; Westoby & Wright, 2003). Size-related traits, such as leaf area (LA), plant height (PHt) and wood anatomical traits (vessel conduit sizes and densities) influence plant competitive ability, aboveground carbon storage, albedo (surface reflectance), and hydrology (Bjorkman et al., 2018; Hietz et al., 2017; Kühn et al., 2021; Plavcová et al., 2024). Quantifying the coordination among traits from different plant parts and diverse functions reveals resource trade-offs and underlying adaptation strategies (Bjorkman et al., 2018; Díaz et al., 2016), while linking environmental factors to plant functional traits is crucial for understanding the impacts of climate change (Bolnick et al., 2011; Dong et al., 2020; Kühn et al., 2021).

While plant strategies based on covariation among leaf morphological traits are well known, those based on twig traits and stem anatomy are mostly unexplored (but see (Chave et al., 2009; Reich, 2014)). Traits like twig increment and diameter are linked to plant architecture, a dimension of plant form that remains underrepresented in trait-based ecology. Yet, architectural traits can offer valuable insights into structure-function relationships in woody species. Recent work (Laurans et al., 2024) has highlighted the ecological importance of such traits, emphasizing their role in linking plant form to function across environmental gradients. Among the anatomical traits, one would expect a plant strategy related to hydraulic safety and conductivity that is expected to be represented by traits such as vessel conduit sizes and densities and tissue fractions in wood (i.e., lignified and parenchymatous). However, current research often lacks an integrated approach that examines the variation in stem, twig, and leaf traits across environmental gradients, particularly those representing species’ central and marginal populations. By integrating data across multiple plant organs and environments, we can more comprehensively understand plant strategies and the impacts of climate change on plants.

Intraspecific trait variation is crucial for plant persistence under environmental changes and constitutes a significant part of community-level trait variation, affecting ecosystem processes like community assembly and nutrient recycling (Bolnick et al., 2011; Jung et al., 2010; Siefert et al., 2015). Although common, studies on intraspecific trait variation across larger geographical gradients are mostly done on tree species and on herbs and rarely on dwarf shrubs (but see Bär et al., 2008; Ropars et al., 2017; Thakur et al., 2024). Consequently, the woody species at the highest elevations in the mountains remain poorly understood despite being vital to alpine ecosystems. Changes in these species can significantly impact ecosystem functioning (Bär et al., 2008; Myers-Smith et al., 2015; Myers-Smith & Hik, 2018). Because they inhabit more extreme climates with high abiotic filtering, biotic factors may play a smaller role in their performance, allowing clearer observation of plant responses to abiotic factors (Paquette & Hargreaves, 2021). This makes them ideal for studying the direct effects of climate on plants.

Plant species have defined geographic distribution limits, and within these ranges, populations can be categorized as either central or marginal based on their position relative to the species’ core distribution area. Studying central and marginal populations is interesting because it reveals how species adapt to optimal and extreme conditions, providing insights into their resilience, evolutionary processes, and responses to climate change (Abeli et al., 2014). In mountainous regions, species are distributed along elevation gradients, with populations at lower elevations marking the warmer range limit, those at higher elevations marking the colder limit, and mid-elevation populations considered central. Similarly, along latitudinal gradients, the northern limit is colder (in the northern hemisphere). Another factor contributing to the formation of species range margins and central populations is water availability that varies greatly in mountains due to rainshadow effect e.g. the northern latitudes in Himalaya. Species spanning these extensive gradients provide opportunities to study trait syndromes in central and marginal populations under specific climates (Halbritter et al., 2018; Thakur et al., 2024), allowing researchers to assess climate change responses across geographically distant populations (Gazol et al., 2015; Thakur et al., 2023). Studies have shown population-specific functional responses to climate and differential climatic adaptations in marginal populations as compared to central populations (Bhuta et al., 2009; Fréjaville et al., 2020; Thakur et al., 2024), emphasizing the need to understand how these populations adapt to changing conditions.

In this study, we investigate radial growth responses and intraspecific trait variations among *Salix flabellaris* populations, a widely distributed Himalayan alpine dwarf shrub. Studying such a alpine high elevation dwarf shrubs is important because they act as sensitive indicators of climate change in mountain ecosystems. Such shrubs are often at the edge of vegetation zones, where environmental conditions are harsh and variable. By examining their adaptations and responses, we can gain insights into the resilience and vulnerability of alpine ecosystems to climate change. Their unique adaptations to extreme conditions also provide valuable information on the limits of plant survival and the potential impacts of global warming on alpine biodiversity.

We analyzed variation in basal area increments, plant age, plant height and 17 stem, twig, and leaf traits in nine populations across wet and warm southern areas, central regions, and, drier and colder northern parts of the Western Himalayas. These populations also spanned elevation gradients from lower to upper species range limits. We, therefore, analyze the intraspecific trait variations of *S. flabellaris* across significant latitudinal and elevational ranges, exploring how they respond to growing season temperature and soil water content. Moreover, we investigate how accelerated warming in Himalayan high mountain regions over the past 20 years has affected the radial growth of *S. flabellaris*, focusing on the context-dependent nature of these effects related to water availability and low-temperature limitation. Finally, we evaluate the implications of significant intraspecific trait variations for predicting the impacts of climate change on the performance and geographic range shifts of *S. flabellaris*.

Specifically, we anticipated that temperature and precipitation gradients defined by both latitudinal and elevational gradients drive growth patterns, longevity and trait variation among *S. flabellaris* populations. Colder temperatures favour conservative strategy (Bjorkman et al., 2018), thus we also expect northern populations (experiencing colder and drier conditions) to exhibit traits favoring conservative life-history strategies. In contrast, southern populations (in warmer and wetter environments) would display traits aligned with more acquisitive strategy. Populations across different latitudes and elevations will likely show distinct trait adaptation syndromes in response to local climatic conditions. We anticipated their shift from acquisitive to conservative life-history strategies as environmental conditions deviate from optima towards colder, drier, or hotter. These variations will be reflected in trait trade-offs, including hydraulic safety versus conductivity, construction cost versus mechanical strength, and enhanced growth versus reduced lifespan. *S. flabellaris* will also exhibit contrasting adaptive responses to variations in growing season temperature depending on water availability, with positive effects in regions of high water availability and negative effects in drier conditions. Finally, we expected a differential radial growth patterns among populations and impact of rising temperatures during the past two decades on radial growth across elevations and latitudes, with northern and high-elevation populations benefiting from reduced temperature constraints. In contrast, southern and mid-elevation populations will be stable due to already favorable conditions. Significant intraspecific trait variations might highlight the necessity for site-specific differentiation when predicting the impacts of climate change on species performance and geographic range shifts.

## METHODOLOGY

### Study Species

This research focuses on *Salix flabellaris* (Figure 1), a dwarf shrub native to the Himalayas. It i an excellent model for studying trait variation due to its extensive distribution across diverse spatial and climatic gradients. As a deciduous broadleaved species, *S. flabellaris* is one of the few woody plants that persist above 4000 meters, defining the climatic shrub line in the region. It begins growing as soon as the snow melts at high elevations and is found in subalpine and alpine regions of the western Himalayas, from 3000 to 4300 meters. Its ability to thrive in extreme climates allows for a clear investigation of abiotic factors with minimal biotic influence. (Mekonnen et al., 2021; Myers-Smith & Hik, 2018). Studying this species’ responses to climat change will enhance our understanding of how alpine woody plants and ecosystems in the Himalayas are affected.

**Figure 1:**
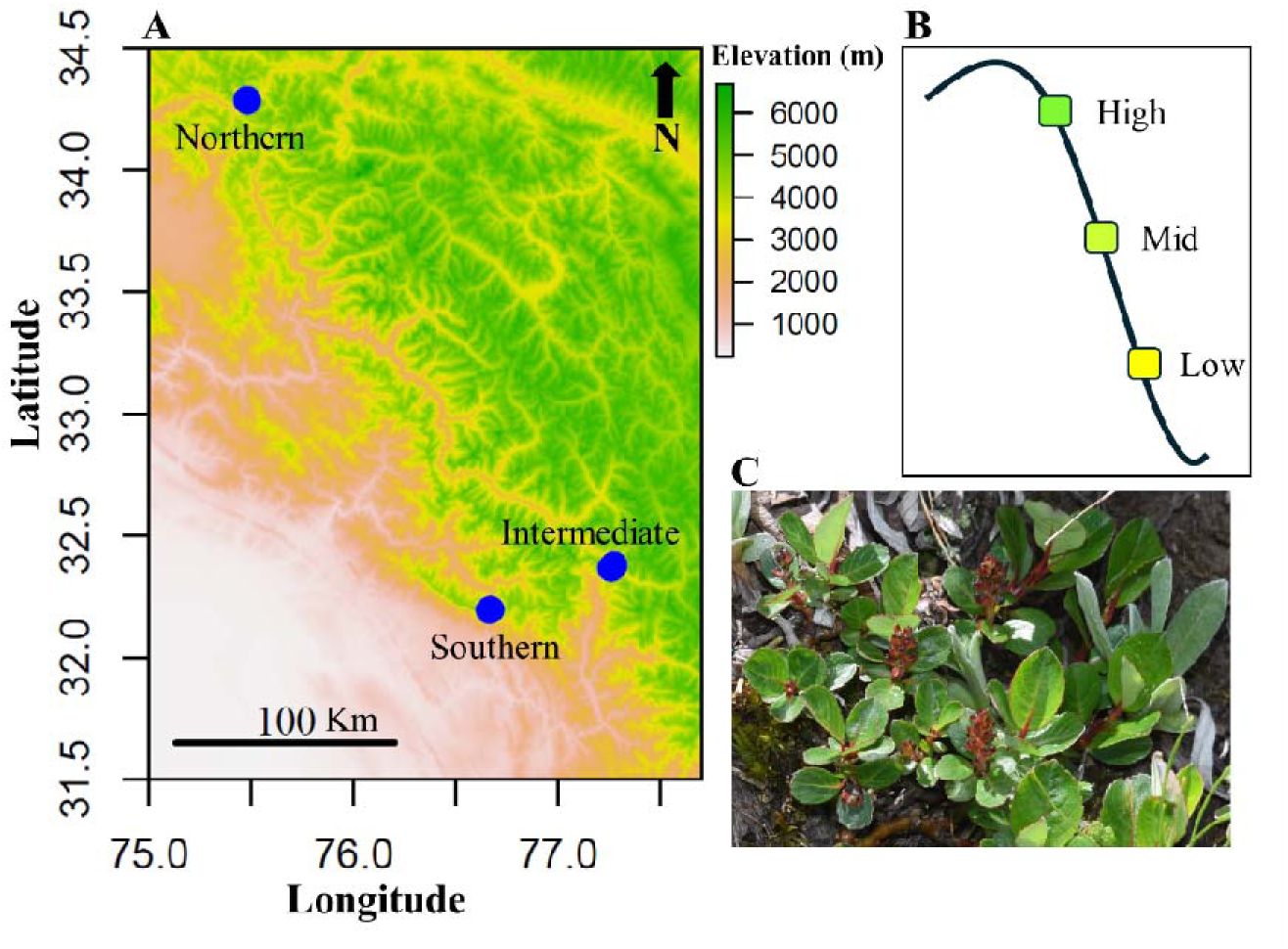
Map showing study area and geographical locations where elevational transects were established (A). In each of the three transects, three elevations (low, mid and high) were studied (represented in B). Northern transect is the coldest and driest while the southern is the wettest and warmest. The part C of the figure represents the focal species. The position of the studied populations in the overall climatic space of the *S. flabellaris* in Western Himalaya is presented in Supplementary Figure S1.

### Study populations and plant sampling

We studied nine populations of the focal species, located between 3200 m and 4200 m above sea level (asl), distributed across three geographically distinct latitudinal transects (Figure 1), with three elevations per transect. The latitudinal transects are labeled northern, intermediate, and southern, while the elevational sites within each transect are classified as low, mid, and high (Figure 1). The northern transect experienced colder temperatures and lower rainfall compared to the others, while the southern transect had the warmest temperatures and highest precipitation levels (Supplementary Table S1 and Figure S2). Consequently, the intermediate transect also experienced intermediate climatic conditions. The mean annual temperature of these populations ranged from 2°C to 5°C, while volumetric soil moisture levels varied from 0.198 to 0.377 based on onsite measurements from 2021 through 2022 using TMS4 dataloggers (Wild et al., 2019).

We collected 76 wood disc samples, with nine samples obtained from each population (except four samples from one of the populations due to the rarity of species at that locality). The samples were collected by cutting a single piece from the thickest stem segment, 3-5 cm long. Within each population, samples were obtained from three plots (3 samples per plot) of approximately 100 m^2^. Within each plot, individuals that were at least 5m apart from each other were sampled. The cut stem samples were immediately placed in a wet paper towel to prevent rapid drying. Within 48 hours of sampling, the stem samples underwent dehydration by being immersed in 50% ethanol for the initial three days, followed by 70% ethanol for the subsequent 7 to 10 days. After the ethanol dehydration process, the samples were air-dried for 72 hours and then stored in paper bags until further processing.

### Growth ring analysis and xylem trait measurements

Plant age and growth data for each of the sampled individual were obtained following established protocols (Doležal et al. 2018). In the laboratory, we utilized a sledge microtome to cut cross-sections from each stem sample. These cross-sections were then stained with Astra Blue and Safranin and permanently affixed to microscope slides using Canada Balsam (Doležal et al. 2022). High-resolution images of the fixed sections were captured using an Olympus BX53 microscope equipped with an Olympus DP73 camera. CellSense Entry 1.9 was employed to analyze the best image obtained from each individual. We measured annual radial growth increments from pith to bark to the nearest micrometer. The age of each individual was estimated by determining the maximum number of annual rings observed in each cross-section. We calculated the basal area increment (BAI) using the growth ring increment data to assess the amount of wood produced and capture changes in annual aboveground productivity. The BAI values were estimated from the inside out (Biondi & Qeadan, 2008) using the ‘bai.out()’ function in the R package dplR (Bunn, 2010).

From the growth rings developed in 2021, we estimated the fraction of lignified tissue (Lig.T), parenchymatous tissue (Par.T), and conductive tissue (Cond.T). These were estimated by randomly selecting a square of 400 um. In addition, the fraction of vessel area (VA), vessel density (V.Den), the size of the largest vessel (V.Max), and the size of the smallest vessel (V.Min) were also estimated.

### Leaf and twig trait measurements

We also measured leaf and twig traits from the same individuals used for wood sampling. The measured leaf traits include leaf area (LA), specific leaf area (SLA), leaf dry matter content (LDMC), and leaf thickness (LT). Leaf traits were measured following recommended protocols (Perez-Harguindeguy et al., 2013; Thakur et al., 2020). The estimated twig traits include twig increment (TL), diameter (TDia), dry matter content (TDMC), specific twig length (STL), twig volume (TVol), and twig density (TDen). Considering that the new twigs may increase in length after the sampling, we measured the twigs formed in the year before the year of sampling. In addition, plant height (PHt) was measured from each sampled individual. These above mentioned traits are related to leaf economics (SLA, LDMC) and twig economics (STL, TDMC, TDen) and plants competitive ability (PHt, LA, TL) to intercept light were selected. TDia, and TVol are also important as twigs with greater diameter or greater volume can accommodate more leaf area that can be beneficial for acquiring more resources.

### Statistical analysis

All ststistical analysis were performed in R software version 4.4.3.

We conducted a principal component analysis (PCA) to examine the coordination between BAI, plant age, plant height, wood anatomy-related traits, leaf traits, and twig traits. While BAI is a growth metric and reflects the outcome of trait × environment interactions rather than intrinsic phenotype, we retained it in the ordination to explore whether certain functional traits are directly associated with plant growth, thereby linking trait variation to performance outcomes. Including BAI allows us to assess whether certain trait combinations are consistently associated with higher or lower growth, thereby providing insight into resource-use strategies and their ecological consequences. To complement the trade-offs and covariation patterns in PCA, we also tested pair-wise correlation among all the studied plant attributes (traits, BAI, plant size and plant age). Correlation analysis serves as a valuable tool to support PCA findings by quantifying the strength and direction of relationships between variables. In particular, it helps to identify trade-offs through negative correlations and reinforces the interpretation of trait covariation patterns that drive the principal components.

To evaluate the influence of climate on all traits collectively, we performed redundancy analysis (RDA) using the rda() function of the vegan package. To ensure that multicollinearity between GST and SWC and with Transect and elevation was not an issue, we calculated variance inflation factors (VIFs), which confirmed that collinearity was low among GST and SWC or Transect and elevation. However, VIFs were higher than 5 when climate (GST and SWC), transect and elevation were together used in RDA model suggesting multicollinearity issue. When only climatic factors or only transect and elevation were used in RDA the VIFs remained below 1.5 suggesting no multicollinearity. Thus we conducted RDA with GST, SWC and their interactions as predictors. Since latitudinal transect and elevation are associated with various local factors beyond climate, we also used separate RDA analysis with latitude and elevation and their interaction as predictors. This analysis was helpful to test mid elevation or mid latitude optima. Subsequently, to get insights into the individual and shared effects of the climatic factors and latitude and elevation, we performed variation partitioning analysis. This was complemented by a series of RDA models designed to quantify the partial and unique variance explained by each factor individually (i.e., of GST, SWC, transect, and elevation). RDA, PCA and variation partitioning analysis were performed using the vegan package in R.

To assess the effects of GST and SWC, and their interaction on each of the estimated plant variable (plant age, plant height and traits), we applied linear mixed effects models, including plot as a random factor. Considering the multicollinearity issue as stated above, similar to the RDA models, we also used separate linear mixed effects models with latitude and elevation and their interaction as predictors. The VIF’s for transect and elevation were also approximately 1. Such additional analysis allowed testing mid latitude or mid elevation optima for each trait. The model structure was as follows: *lmer(response ∼ predictor1 * predictor2+(1|Plot), data = data)*.

Although the absolute elevation values differed among transects, we treated elevation as a categorical variable (low, mid, high) in RDA and mixed effect models to reflect relative positions along the local elevational gradient within each transect. This approach allowed us to compare growth responses across consistent ecological zones (e.g., lower, mid, and upper limits of species occurrence in studied transects) rather than relying on absolute elevation, which may not be ecologically equivalent across regions due to differences in climate, vegetation, and site conditions.

To analyze differences in basal area increments (BAI) among nine populations and explore changes over the last two decades (since the year 2000) in each of the population, we again used a linear mixed effects model, testing the effects of growth year, latitudinal transect (three elevational transects at different locations), elevation (three elevations within each transect), and their interactions on BAI with plot as a random factor. Plant age (log-transformed) was included as a covariate, as it can significantly influence growth in woody species. The model in this case was as follows: *lmer(log(BAI)∼log(Age)+(Year of growth+Transect+Elevation)^2+(1|Plot), data = data)*.

Log-transformed age was used because preliminary comparisons of models testing age, sqrt(age), and log(age) indicated that log(age) was the best predictor of BAI, likely due to a greater increase in BAI during the early stages of plant development (Bigler, 2016; Dolezal et al., 2021; Thakur et al., 2024). In this study, plant age referred to the age at the time BAI was estimated rather than the final age of the individual plants. We limited the BAI analysis to the past two decades to ensure sufficient data from enough individuals per population. This approach allowed us to explore how annual productivity, measured by BAI, has changed over the last 20 years. Because GST and SWC data were only available for a single year, we could not run alternate models including these climatic variables in the BAI model. BAI was log-transformed before fitting the models to ensure normality.

## RESULTS

### Independent dimensions of variation in stem, twig and leaf traits

The principal component analysis (PCA) of all populations revealed independent variation among stem, twig, and leaf traits. The first five axes accounted for 64.8% of the variation in the multi-dimensional functional space, with individual axes explaining between 19.8% and 8.4%. The first two axes together explained 32.2% of the variation (Figure 2A). The first axis was primarily associated with traits related to xylem structure and function, including lignified, conductive, and parenchymatous tissue, as well as vessel size. The second axis was linked to twig construction cost and strength traits, such as STL, twig diameter, and twig volume. The third axis explained 12.5% of the variation and captured the leaf economic spectrum, represented by LDMC and SLA (Supplementary Figure S3). The fourth axis, explaining 9.7%, was related to water transport traits, particularly vessel size and density. The fifth axis, which explained 8.4%, represented an independent dimension of variation involving plant height and its correlation with twig increment (Supplementary Figure S3). These trait dimensions were further supported by correlation analyses (Supplementary Figure S4), which revealed negative correlations among traits involved in trade-offs.

**Figure 2:**
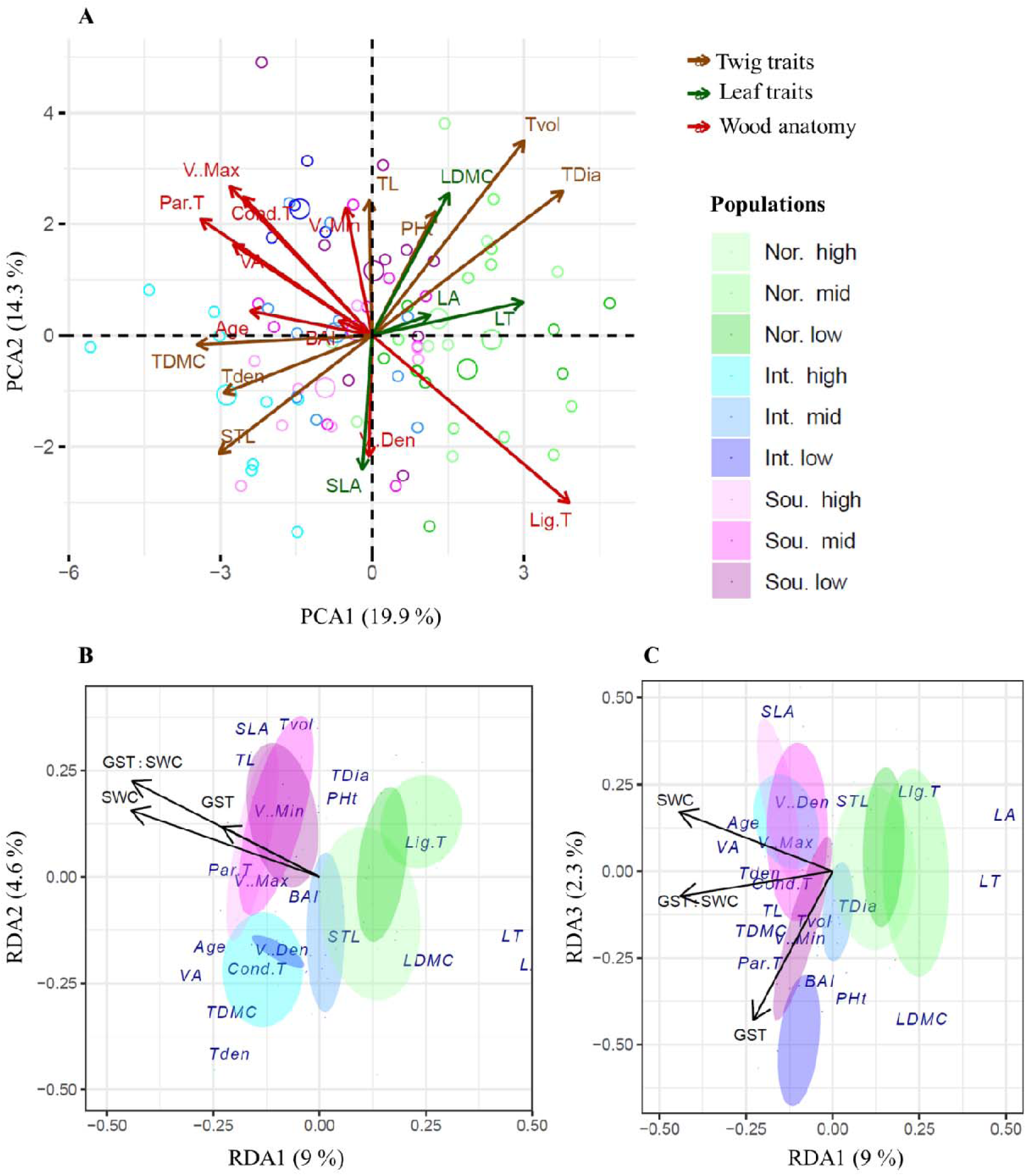
Principal component analysis (PCA) biplot showing the covariation of various traits in multivariate space (A) and redundancy analysis plots (B and C) showing the effect GST, SWC and their interaction on all studied traits along the first three axes. Plot B includes axis 1 versus axis 2 and the plot C presents axis 1 versus axis 3. Bigger circles in Figure A represent population centroids and maller ones represent individual in a population. Shaded areas in plot B and C represent 50% confidence intervals around the centroid of each of the population. In population codes, Int. = Intermediate; Nor. = Northern; and Sou. = Southern latitudinal transect and high, mid and low are the three elevations in each latitudinal transect.

### Trait variation in multivariate trait space

The redundancy analysis (RDA) using climate variables as predictors revealed a significant effect of growing season temperature (GST), soil water content (SWC), and their interaction, explaining a total of 15.85% of the variation in traits (Figure 2B and 2C). SWC had the strongest influence, accounting for 6.9% of the variation, followed by its interaction with GST (5.15%) and GST alone (3.82%). Traits associated with higher SWC and GST included lower values of lignified tissue, LDMC, leaf area, and leaf thickness, along with higher values of plant age, vessel area, parenchymatous tissue, twig density, and TDMC. Traits linked to increased GST included lower STL, SLA, and leaf area, greater BAI, LDMC, plant height, and the fraction of parenchymatous tissue in the stem (Figure 2B and 2C).

In the RDA with latitudinal transect and elevation as predictors, both predictors together explained 32.07% of the variation in the traits studied. The first two axes accounted for 15.1%, and 7.15% of the total variation, respectively (Supplementary Figure S6). Both predictors and their interaction had significant effects on traits. The latitudinal transects had the strongest influence (p < 0.001), explaining 19.9% of the variation, followed by its interaction with elevation (p < 0.006), which accounted for 6.87%, and then elevation alone (p < 0.001), explaining 5.27% of the variation. Populations from the northern transect were separated from the others along the first axis, while populations from the southern and intermediate transects were distinguished along the second axis. Traits associated with northern populations included larger leaf area (LA), leaf thickness (LT), leaf dry matter content (LDMC), plant height (PHt), twig diameter, and a higher fraction of lignified tissue in the stem (Supplementary Figure S6). In contrast, southern populations exhibited traits such as higher specific leaf area (SLA), greater twig length increment, and larger vessel area. Populations from the intermediate transect were characterized by traits like higher twig dry matter content (TDMC), twig density, stem twig length (STL), and a greater fraction of parenchymatous tissue in the stem (Supplementary Figure S6).

In the variance partitioning analysis, elevation and SWC did not explain significant unique variation in plant traits, instead, their effects were largely shared with other factors. In contrast, transect explained a substantial amount of unique variation, with more than half of its contribution being independent of climate and elevation (Figure S5). This suggests that non-climatic factors associated with transect, such as biogeographic or historical influences, play an important role in shaping trait variability. Similarly, a portion of the variation attributed to GST remained unique, indicating a distinct climatic influence.

### Effect of climatic factors on plant traits

Plant age and two stem anatomy-related traits were significantly influenced by climate (Table 1, Figures 3). Age decreased with increasing temperature at higher soil water content (SWC), but the trend reversed at lower SWC (Figure 3A). The fraction of conductive tissue in the stem and vessel area were affected by the interaction between growing season temperature (GST) and SWC (Table 1). Both parameters tended to increase with temperature at lower soil moisture levels but showed the opposite pattern at higher moisture content (Figures 3 B-C).

**Figure 3:**
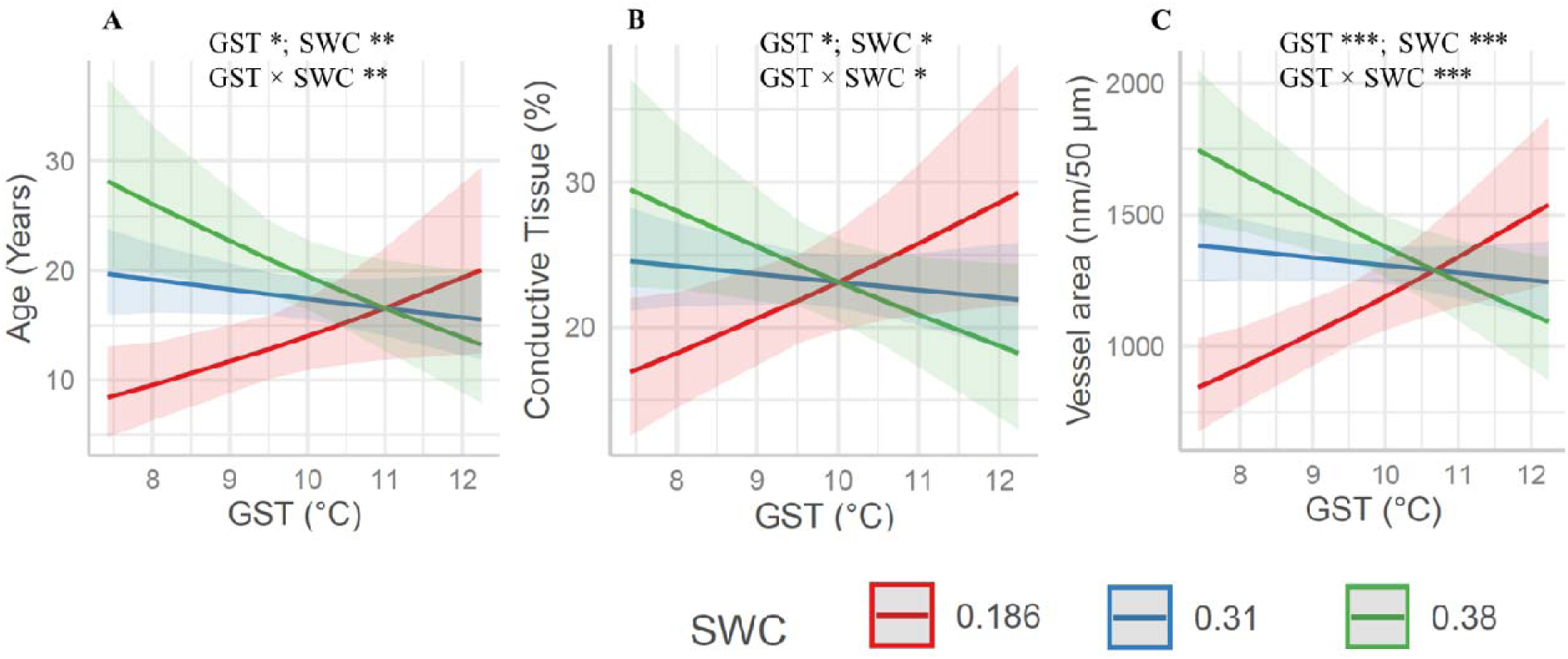
Plots showing the changes in plant age and other wood anatomical variables with temperature (GST) and/or soil water content (SWC). Figures are based on model statistics shown in supplementary table 2. Only traits with significant effects of any of the factors are shown here. Low, mid and high precipitation levels are plotted to avoid complexity in figures. Shaded area represent 95% confidence intervals. n.s. = non significant; * = p<0.05; ** = p<0.01; *** = p<0.001.

**Table 1:** Summary of the analyses investigating the effect of growing season temperature (GST), soil water content (SWC) and their interactions on different variables estimated from *Salix flabellaris*. P value in bold represents significant effects.

| Trait type | Trait | Abbreviation | GST |  | SWC |  | GST × SWC |  |
| --- | --- | --- | --- | --- | --- | --- | --- | --- |
|  |  |  | F | P | F | P | F | P |
| Age and stem anatomy related | Plant age (years) | Age | 6.304 | **0.014** | 8.435 | **0.005** | 6.633 | **0.012** |
|  | % lignified tissue | Lig. T | 3.503 | 0.074 | 2.72 | 0.113 | 2.297 | 0.144 |
|  | % parenchymatous tissue | Par. T | 0.813 | 0.377 | 0.373 | 0.547 | 0.252 | 0.62 |
|  | % conductive tissue | Cond. T | 5.84 | **0.018** | 6.12 | **0.016** | 5.809 | **0.019** |
|  | Vessel density (in 50 µm) | V. Den | 2.745 | 0.111 | 3.944 | 0.059 | 3.544 | 0.072 |
|  | Vessel area (nm in 50 µm) | VA | 12.497 | **0.001** | 14.670 | **0.001** | 12.26 | **0.001** |
|  | Minimum vessel size (nm) | V. Min | 0.018 | 0.895 | 0.153 | 0.697 | 0.209 | 0.649 |
|  | Maximum vessel size (nm) | V. Max | 0.382 | 0.538 | 0.586 | 0.446 | 0.344 | 0.559 |
| Height and twig related | Plant height (cm) | PHt | 1.272 | 0.263 | 2.989 | 0.088 | 2.601 | 0.111 |
|  | Twig length increment (mm) | TL | 0.979 | 0.326 | 1.353 | 0.249 | 1.961 | 0.166 |
|  | Twig volume increment (mm<sup>3</sup>) | TVol | 4.578 | **0.043** | 5.877 | **0.024** | 6.406 | **0.019** |
|  | Twig diameter (mm) | TDia | 4.248 | **0.05** | 5.582 | **0.027** | 5.388 | **0.029** |
|  | Twig density | TDen | 21.314 | **<0.001** | 22.834 | **<0.001** | 21.616 | **<0.001** |
|  | Twig dry matter content (g/g) | TDMC | 5.691 | **0.026** | 5.182 | **0.033** | 4.500 | **0.036** |
|  | Specific twig length (mm/g) | STL | 0.004 | 0.95 | 0.113 | 0.74 | 0.119 | 0.733 |
| Leaf | Leaf dry matter content (g/g) | LDMC | 0.626 | 0.438 | 0.034 | 0.856 | 0.375 | 0.547 |
|  | Specific leaf area (cm<sup>2</sup>/g) | SLA | 6.441 | **0.019** | 3.537 | 0.073 | 5.183 | **0.033** |
|  | Leaf area (cm<sup>2</sup>) | LA | 0.488 | 0.491 | 0.583 | 0.453 | 0.006 | 0.936 |
|  | Leaf thickness (mm) | LT | 0.797 | 0.381 | 1.085 | 0.309 | 0.345 | 0.560 |

Among twig traits (Figure 4 A-D), twig length increment, volume, and diameter exhibited similar variation patterns with GST and SWC (Table 1). These traits increased with rising GST at higher moisture levels but decreased at lower SWC. Twig density and twig dry matter content (TDMC) showed similar trends to each other, but their patterns were opposite to that of twig diameter. For leaf traits, climate significantly affected only specific leaf area (SLA) (Table 1 and Figure 4 E), which decreased with increasing GST at lower SWC but increased at higher SWC. SLA was also lower under drier conditions.

**Figure 4:**
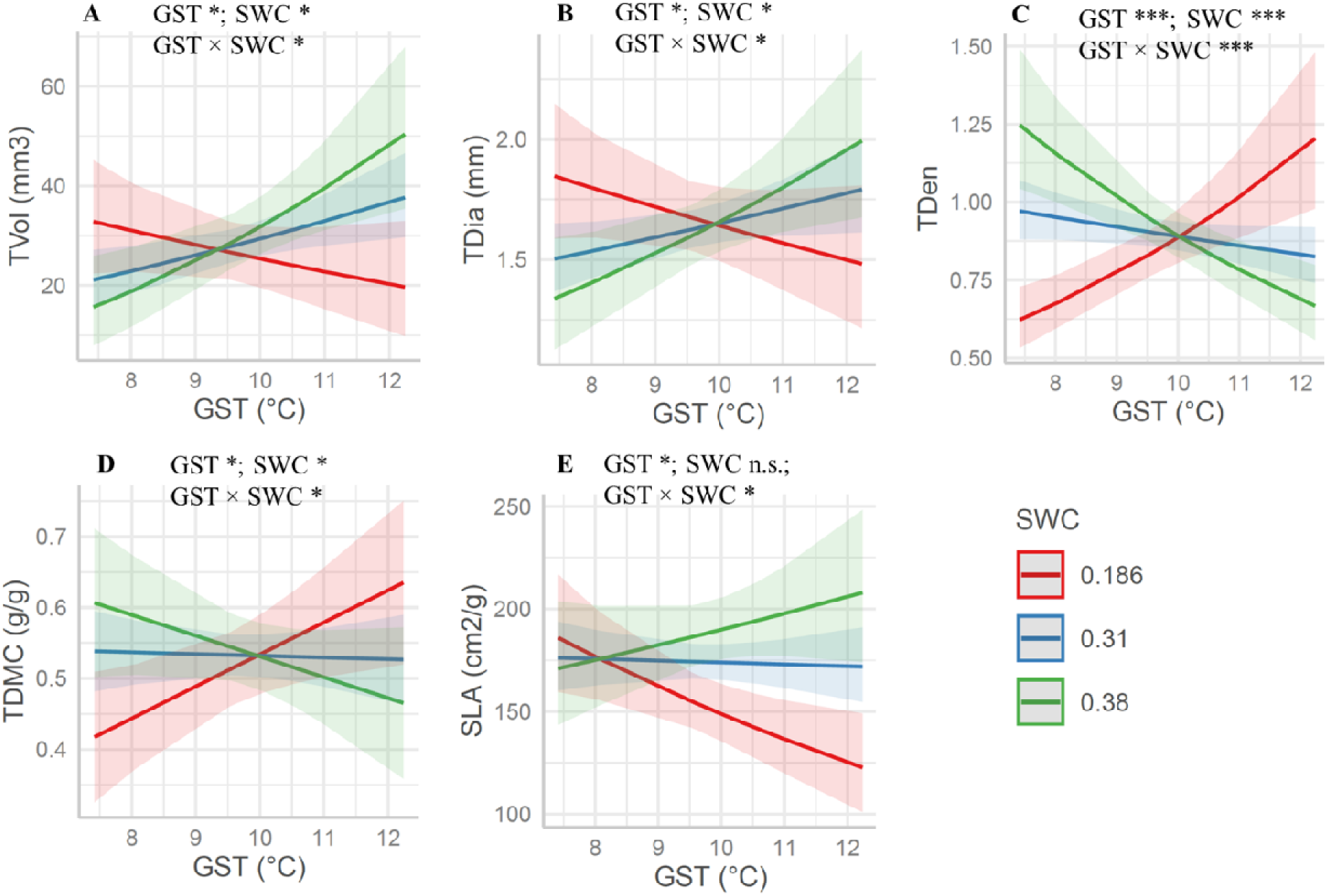
Plots showing the changes in plant height, twig traits and leaf traits with temperature (GST) and/or soil water content (SWC). Figures are based on model statistics shown in supplementary table 2. Only traits with significant effects of any of the factors are shown here. Low, mid and high precipitation levels are plotted to avoid complexity in figures. Shaded area represent 95% confidence intervals. n.s. = non significant; * = p<0.05; ** = p<0.01; *** = p<0.001.

### Effect of latitudinal and elevational sites on each of the variable

With the exceptions of plant age, vessel size, vessel density, and conductive tissue, all the studied traits were significantly influenced by at least one factor i.e., latitudinal transect, elevation, or their interaction (see Table 2 reporting linear model statistics).

**Table 2:**
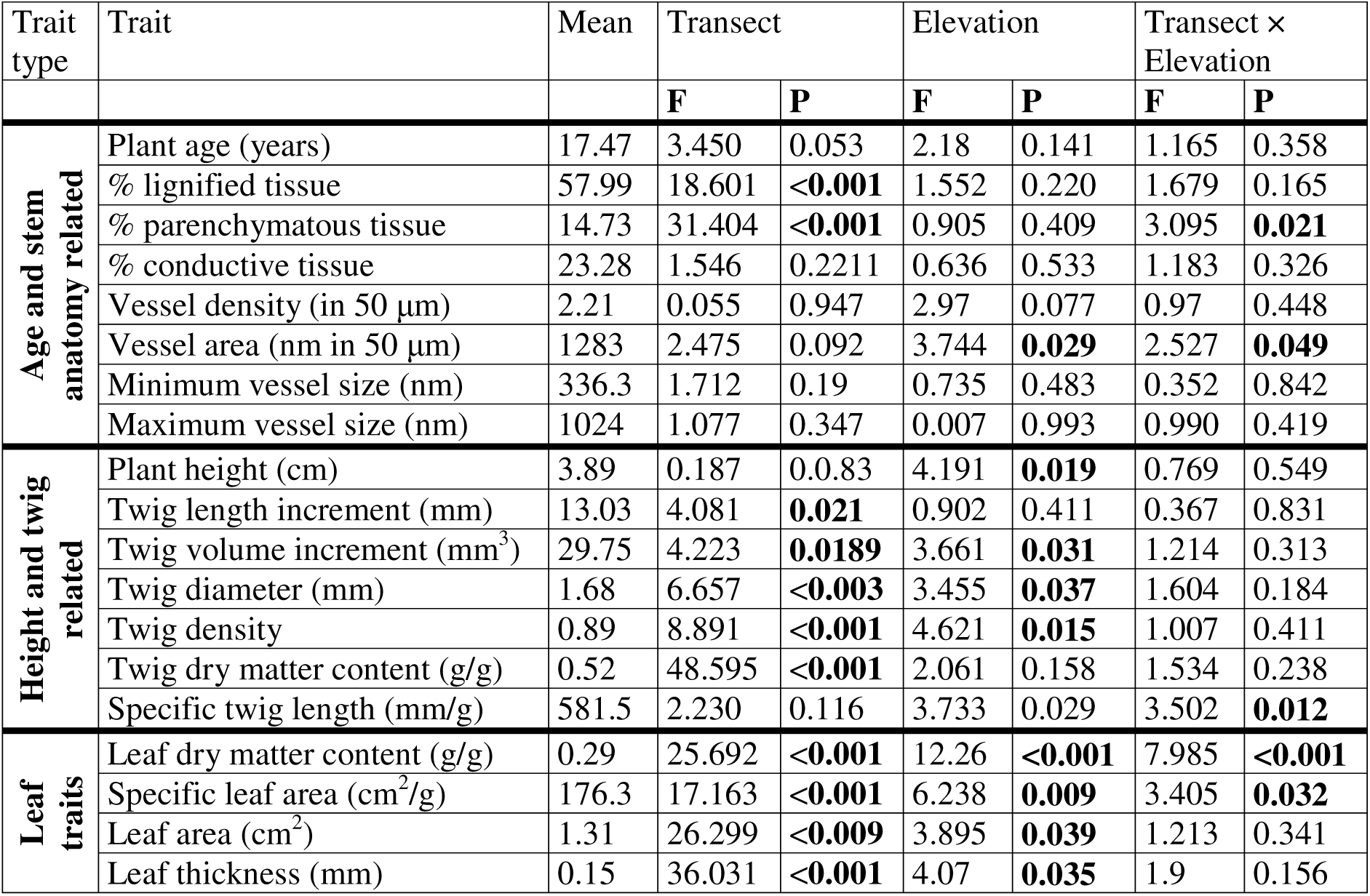
Summary of the analyses investigating the effect of transect, elevation and their interaction different functional traits estimated from *Salix flabellaris*. P value in bold represents significant effects.

| Trait type | Trait | Mean | Transect |  | Elevation |  | Transect × Elevation |  |
| --- | --- | --- | --- | --- | --- | --- | --- | --- |
|  |  |  | F | P | F | P | F | P |
| Age and stem anatomy related | Plant age (years) | 17.47 | 3.450 | 0.053 | 2.18 | 0.141 | 1.165 | 0.358 |
|  | % lignified tissue | 57.99 | 18.601 | <**0.001** | 1.552 | 0.220 | 1.679 | 0.165 |
|  | % parenchymatous tissue | 14.73 | 31.404 | <**0.001** | 0.905 | 0.409 | 3.095 | **0.021** |
|  | % conductive tissue | 23.28 | 1.546 | 0.2211 | 0.636 | 0.533 | 1.183 | 0.326 |
|  | Vessel density (in 50 µm) | 2.21 | 0.055 | 0.947 | 2.97 | 0.077 | 0.97 | 0.448 |
|  | Vessel area (nm in 50 µm) | 1283 | 2.475 | 0.092 | 3.744 | **0.029** | 2.527 | **0.049** |
|  | Minimum vessel size (nm) | 336.3 | 1.712 | 0.19 | 0.735 | 0.483 | 0.352 | 0.842 |
|  | Maximum vessel size (nm) | 1024 | 1.077 | 0.347 | 0.007 | 0.993 | 0.990 | 0.419 |
| Height and twig related | Plant height (cm) | 3.89 | 0.187 | 0.083 | 4.191 | **0.019** | 0.769 | 0.549 |
|  | Twig length increment (mm) | 13.03 | 4.081 | **0.021** | 0.902 | 0.411 | 0.367 | 0.831 |
|  | Twig volume increment (mm<sup>3</sup>) | 29.75 | 4.223 | **0.0189** | 3.661 | **0.031** | 1.214 | 0.313 |
|  | Twig diameter (mm) | 1.68 | 6.657 | <**0.003** | 3.455 | **0.037** | 1.604 | 0.184 |
|  | Twig density | 0.89 | 8.891 | <**0.001** | 4.621 | **0.015** | 1.007 | 0.411 |
|  | Twig dry matter content (g/g) | 0.52 | 48.595 | <**0.001** | 2.061 | 0.158 | 1.534 | 0.238 |
|  | Specific twig length (mm/g) | 581.5 | 2.230 | 0.116 | 3.733 | 0.029 | 3.502 | **0.012** |
| Leaf traits | Leaf dry matter content (g/g) | 0.29 | 25.692 | <**0.001** | 12.26 | <**0.001** | 7.985 | <**0.001** |
|  | Specific leaf area (cm<sup>2</sup>/g) | 176.3 | 17.163 | <**0.001** | 6.238 | **0.009** | 3.405 | **0.032** |
|  | Leaf area (cm<sup>2</sup>) | 1.31 | 26.299 | <**0.009** | 3.895 | **0.039** | 1.213 | 0.341 |
|  | Leaf thickness (mm) | 0.15 | 36.031 | <**0.001** | 4.07 | **0.035** | 1.9 | 0.156 |

Lignified tissue in the stem was lowest at the intermediate transect and highest at the northern transect (Figure 5A). In contrast, the fraction of parenchymatous tissue showed an opposite pattern, with the highest values at the intermediate transect. However, elevational patterns for parenchymatous tissue were inconsistent across transects and elevations, being lower at mid-elevations in the intermediate transect, while the opposite was observed in the northern transect (Figure 5B). Vessel area was higher at higher elevations (Figure 5C) and decreased from the southern to northern transects at mid-elevation, with no change at other elevations.

**Figure 5:**
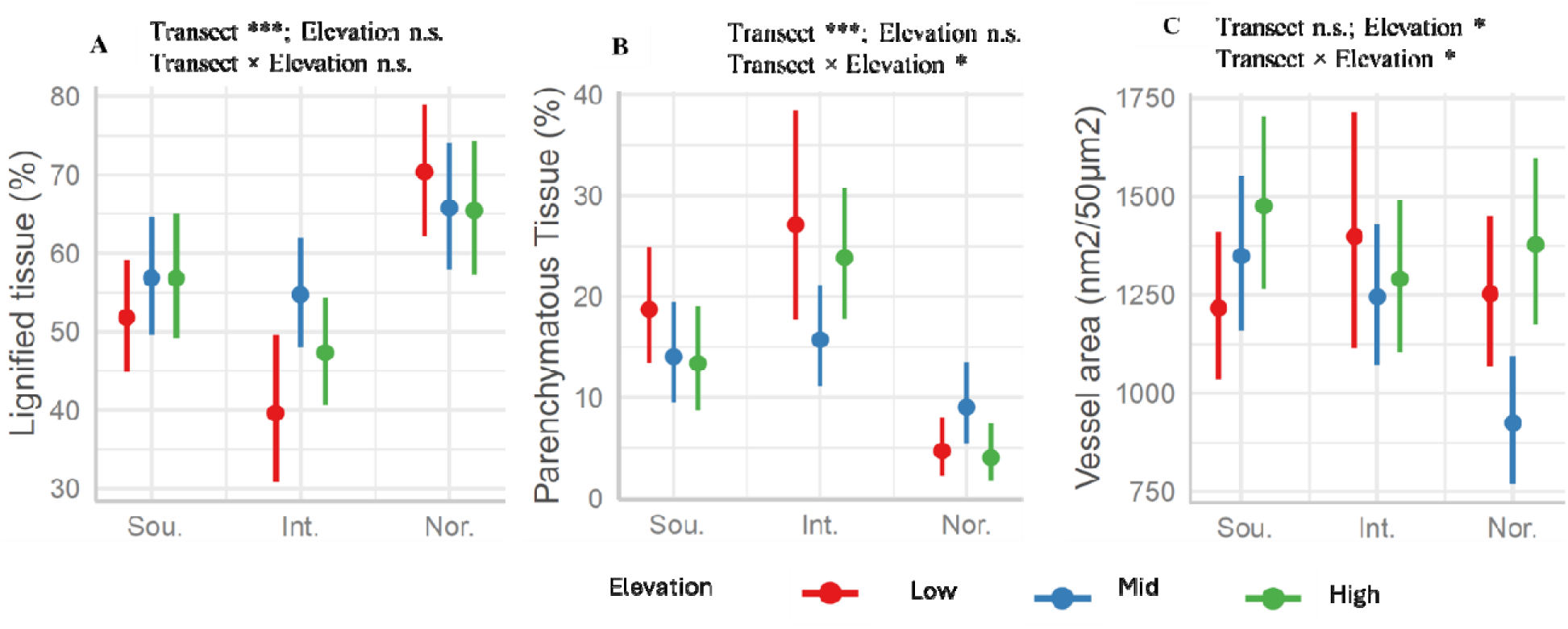
Differences in stem wood anatomy among populations at three different elevations of each of the three latitudes. Variables without any significant effect are not shown. Error bar represent 95% confidence intervals. n.s. = non significant; * = p<0.05; ** = p<0.01; *** = p<0.001.

For plant height and twig traits, latitudinal transects and elevation had strong individual effects, with an interactive effect observed only for stem twig length (STL) (Table 2, Figure 5 A-G). Plant height was solely affected by elevation, with taller plants found at lower elevations and shorter plants at higher elevations. Twig length increment was greatest in populations from the southern transect and smallest in the northern transect. Twig volume increment was highest in the northern transect and lowest in the intermediate transect, with a general decrease at higher elevations. Twig density followed a similar pattern, being highest in the northern transect, lowest in the intermediate transect, and declining with elevation. Both twig density and TDMC exhibited similar trends, with higher values in the intermediate transect. Among elevations, twig density was lower at mid-elevations. Twig construction cost (high STL = low cost) generally decreased with increasing elevation, except in the northern transect, where it was higher at mid-elevation.

Leaf traits, particularly those related to resource use strategy (SLA and LDMC), were most significantly affected (Table 2, Figure 6 H - I), with all effects (both individual and interactive) being significant. LDMC was highest at low elevations in the intermediate transect and lowest at mid and high elevations in the southern transect. In the northern transect, LDMC was lowest at low elevation, whereas the opposite pattern was observed in the other two transects. SLA was highest in the southern transect and decreased with elevation, except in the northern transect, where mid-elevation exhibited higher values than the other two elevations. Leaf area was lowest in the southern transect and highest in the northern transect, with mid-elevations showing th greatest leaf area overall. Leaf thickness followed a similar pattern to leaf area, except in th northern transect, where it increased with elevation.

**Figure 6:**
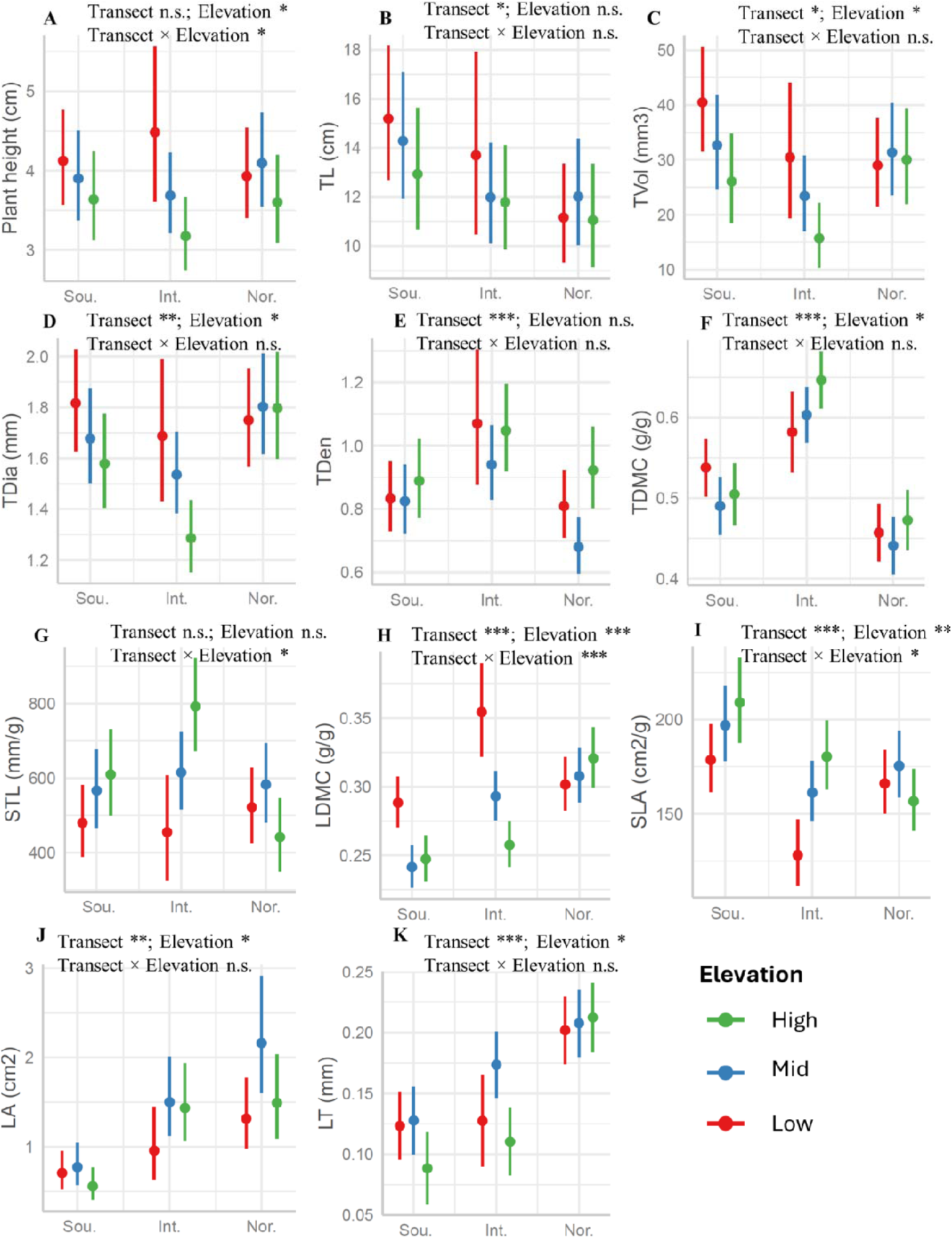
Differences in plant height (A), twig traits (B-G) and leaf traits (H-K) among populations at three different elevations of each of the three transects. Error bars represent 95% confidence intervals.

### Differences in basal area increment (BAI) and temporal trends in BAI

Basal area increment (BAI) was highest at mid-elevations, a pattern consistent across latitudinal transects (Table 3, Figure 7A) suggesting optimal growth at mid elevations. The effect of growth year on basal area increment (BAI) indicates a significant increase from 2000 to 2021 (Figure 7B). Additionally, the growth year significantly interacted with elevation (Supplementary Table 2), suggesting that changes in BAI over the past two decades have been inconsistent across elevations (Table 3 and Figure 7B). The increase occurred only at high elevations, while low and mid-elevations showed no significant change (Figure 7B). There was no significant interaction between the year of growth and latitudinal transect, indicating that BAI changes over the past two decades were similar across latitudinal transects.

**Figure 7:**
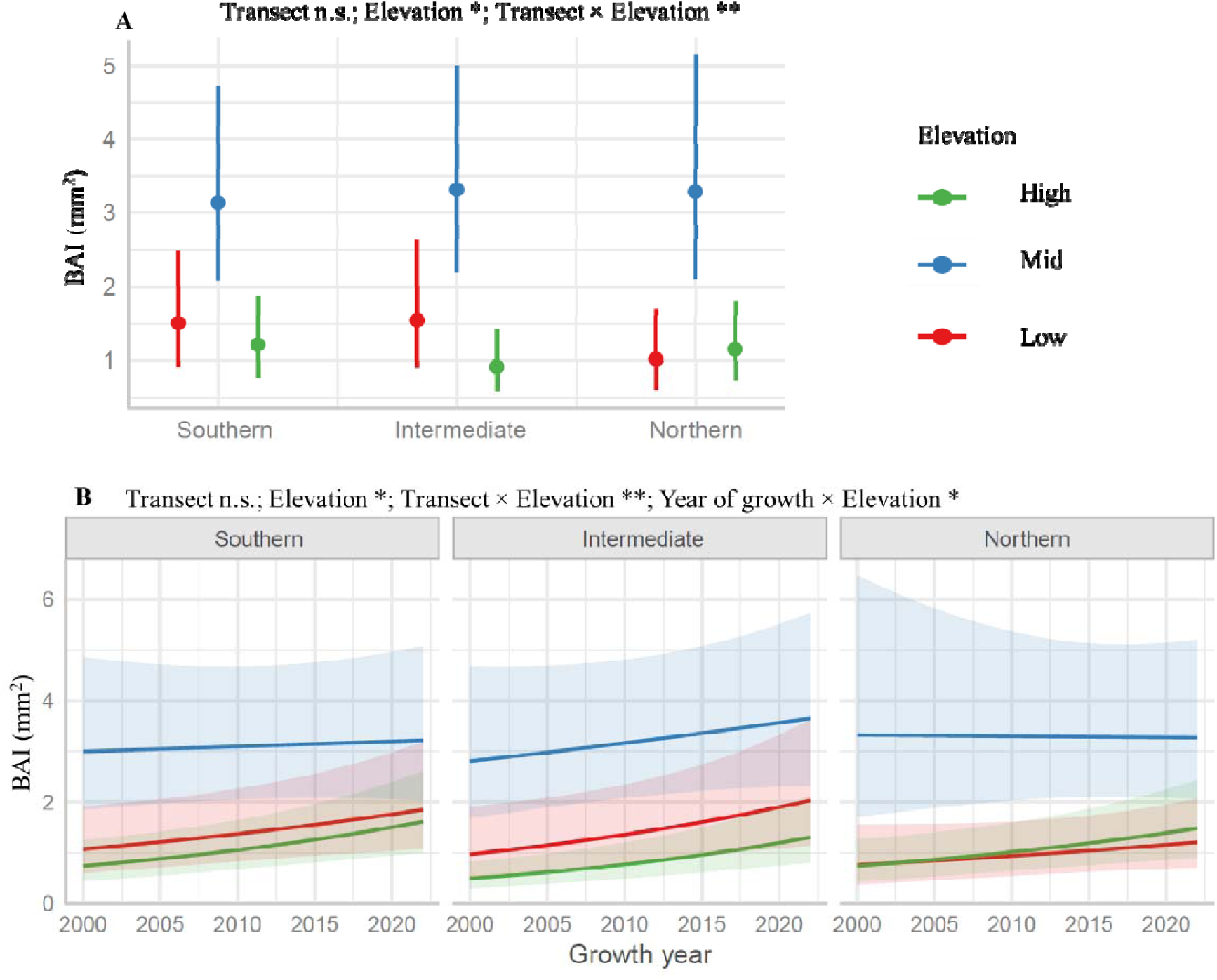
Differences in basal area increment (BAI) among populations (A) and changes in it during last two decades across populations (B). Populations were at three different elevations at each of the three latitudinal transects (northern, intermediate, southern; also see Figure 1). The shaded areas represent 95% confidence intervals. The change in growth from 2000 to 2021 wa significant only at high elevations. n.s. = non significant; * = p<0.05; ** = p<0.01.

**Table 3:** Linear mixed effects model statistics showing the effect of transect and elevation on BAI for the last two decades of growth. P value in bold represents significant effects.

| Predictors | F value | P value |
| --- | --- | --- |
| Plant age (covariate) | 176.1 | **<0.001** |
| Transect | 0.5 | 0.619 |
| Elevation | 3.9 | **0.021** |
| Year of growth | 12.3 | **<0.001** |
| Transect × Elevation | 3.7 | **0.005** |
| Transect × Year of growth | 0.5 | 0.62 |
| Elevation × Year of growth | 3.8 | **0.023** |

## DISCUSSION

We reported independent plant strategies as represented by stem anatomy, twig traits and leaf traits. Growing season temperature and soil water content significantly influenced stem anatomy, and twig and leaf traits. Higher temperatures improved shrub performance, including larger twigs and lower construction costs, but only when soil water availability was high. Mid-elevation populations exhibited optimal performance across latitudes, characterized by higher stem growth rates, lower twig construction costs, less dense twigs, and larger leaves. Northern populations showed more lignified stems, thicker leaves, greater twig diameter and higher leaf dry matter content. From 2000 to 2021, stem growth (basal area increment) increased with rising temperatures, benefiting high-elevation populations the most, while mid-elevation populations remained stable possibly due to already favorable conditions.

### Stem, twig and leaf traits vary largely independently of each other

This study shows that stem, twig and leaf traits vary largely independently of each other and thus represent different plant adaptation strategies. Based on structural connectivity to one another (the xylem in the stem is connected to twigs, which in turn bear leaves), the expectation was the presence of strong covariation among the studied traits.

The primary dimension of trait variation associated with stem wood anatomy was linked to the hydraulic safety-conductivity strategy, reflecting a trade-off between lignified parenchymatous tissue and conduit vessel tissues. In this trade-off, increasing hydraulic safety leads to a decrease in the ability to conduct water. Another trade-off involved vessel density and minimum vessel area. Although we anticipated this trade-off to align with the hydraulic safety-conductivity dimension, its independence suggests that hydraulic functions in plants are much more complex and regulated through multiple trade-offs. This complexity highlights the significance of these traits for plant survival under changing climatic conditions. While many previous studies have reported trade-offs among vessel size and density (Hietz et al., 2017), our findings reveal that maximum vessel size is traded off against lignified tissue, while minimum vessel size is traded off against vessel density. This suggests that minimum and maximum vessel sizes reflect two distinct strategies in the studied species.

The independent strategy was associated with twig traits. It was represented by specific twig length and diameter, being parallel to findings from previous studies on root traits, particularly the trade-off between specific root length and root diameter (Bergmann et al., 2017; Carmona et al., 2021; Münzbergová et al., 2025). However, since we did not examine root traits in this study, the relationship between twig and root traits remains unclear. In roots, diameter is often linked to collaboration with symbiotic partners, while in twigs, diameter primarily relates to accommodating more conductive tissue and providing greater mechanical strength. This dimension balances construction costs with mechanical strength in twigs. In leaves, the trade-off aligns with the well-known exploitative-conservative strategy (Wright et al., 2004), where specific leaf area (SLA) and leaf dry matter content (LDMC) trade off with each other. Overall, stem anatomy, twig, and leaf traits are functionally connected, yet they largely vary independently and represent different intraspecific plant strategies.

### Warming effects on plant functions depend on water availability

Apart from trait variation along latitudinal and elevational gradients, we also examined how climate influences various plant traits. Our observations revealed distinct patterns related to plant age, in response to temperature and soil moisture levels. Specifically, plant age tends to increase with increasing growing season temperature at sites with lower soil moisture, but this trend reverses at sites with higher moisture levels. This pattern may be attributed to the increased likelihood of frost-related plant mortality in drier and colder conditions, particularly during winter, when plants are more exposed to cold air (Dolezal et al., 2016). In contrast, at wetter sites with warmer climates, plant age may be influenced by greater biotic stress, as well as higher growth rates associated with these conditions. Previous studies have shown that rapid growth in such environments often correlates with shorter plant lifespans (Bigler, 2016; Dolezal et al., 2021; Thakur et al., 2024). Furthermore, an increase in fraction of conductive tissue and vessel area with rising temperatures under conditions of lower water availability could be because higher temperatures elevate water demand, and an increase in these traits can enhance plant’s ability to meet increased demand for water.

Among twig and leaf traits, twig traits demonstrated a stronger correlation with climate than leaf traits. Since twig traits are less frequently measured, it is difficult to compare these results with previous studies. Low temperatures are a key factor restricting plant growth at high elevations (Körner, 2003). In this context, the observed increases in twig volume and diameter with rising temperatures were as expected. At higher soil moisture levels, the growth of thicker but less dense twigs may be attributed to more favorable environmental conditions, allowing plants to produce twigs that can support a greater number of leaves with less investment. Conversely, the combination of higher temperatures and lower water availability creates stressful conditions, likely explaining the decrease in twig diameter and volume under drier conditions. Additionally, the increase in twig density and twig dry matter content with rising temperatures at lower water availability may reflect a response to the increased demand for water in drier, warmer environments. Plants may develop tougher, denser tissues to mitigate the risk of cavitation in such conditions, which necessitate a greater investment in dry matter.

SLA generally increases with temperature (Rosbakh et al., 2015; Wilcox et al., 2021). However, our findings showed a more nuanced pattern of interacting effects of temperature and moisture as also found in (Kosová et al., 2022). SLA decreased with rising temperatures at lower moisture levels but increased at higher moisture levels. This divergence under drier conditions could be due to the combination of higher temperatures and lower water availability, which creates dehydrating conditions (because of increased evapotranspiration). Under such stress, plants likely invest more in each unit of leaf area, possibly by allocating resources to thicker cell walls and conductive tissues to enhance water retention and reduce the risk of dehydration.

While latitudinal transects, elevation, and climate significantly influenced most of the studied traits, it is crucial to recognize that trait variation can also stem from genetic differences among populations and other factors such as soil properties and competition at both intraspecific and interspecific levels. These factors were not included in this study. Acknowledging these additional influences is important, as they can provide a more comprehensive understanding of plant strategies and responses to climate change. By separating the effects of these factors from climatic influences, future research could yield valuable insights into how plants adapt and thrive in varying environmental conditions.

### Central populations are functionally distinct from marginal populations

Latitude and elevation significantly influenced the functioning of studied populations, affecting their fitness and long-term persistence. Among various traits, the basal area increment (BAI), which indicates plant growth and productivity, was notably higher at mid-elevations across all three latitudinal transects. Central populations at mid-elevations also had traits suggesting better performance than marginal populations at lower and upper elevations. This enhanced performance at mid-elevations is likely due to an optimal temperature. While other abiotic factors such as nutrient availability and wind exposure also vary with elevation, their role in enhancing growth is unlikely as nutrient availability generally decreases and wind exposure increases with elevation, both of which tend to limit rather than promote plant performance. Herbivory and pathogens may reduce growth at lower, warmer elevations, while low temperatures at higher elevations generally limit plant growth (Körner, 2003; Paquette & Hargreaves, 2021). Additionally, traits such as leaf area and leaf thickness, associated with efficient solar energy use, peaked at mid-elevations. Among twig traits, twig density was lower at mid-elevations, reflecting reduced carbon investment per unit volume.

The intermediate transect, representing non-marginal populations along a latitudinal gradient (also intermediate in terms of climatic conditions), displayed several distinct traits indicative of specific adaptive strategies. Notably, these populations showed an increased proportion of parenchymatous tissue and a reduced fraction of lignified tissue in stem wood. This tissue composition, along with a lower twig diameter, twig volume, and higher specific twig length, suggests lower investment in supportive tissues. This reduction in lignified tissue and the smaller twig dimensions and lower twig construction cost per unit length may reflect the less extreme environmental conditions at the central transect, where there is a balance between temperature and water availability, reducing the need for robust supportive structures.

The intermediate transect populations exhibited higher twig density and twig dry matter content (TDMC). These traits typically relate to higher investment in tissue construction. However, when combined with specific twig length (STL) patterns, it implies a strategic allocation of resources that allows the plants to form thinner, denser twigs that can bear enough leaves and are also strong enough to withstand environmental fluctuations. The lower specific leaf area (SLA) observed in these populations indicates higher leaf construction costs, which might initially seem disadvantageous. However, a lower SLA is often associated with a more conservative resource-use strategy, enabling plants to maintain their resources over an extended period (Wright et al., 2004). This trait can be particularly beneficial throughout the growing season, as it allows for more efficient resource utilization, ensuring sustained growth and resilience. While SLA alone cannot fully capture the complexity of resource-use efficiency or resilience, it provides valuable insight into a plant’s adaptive strategy, particularly in environments where sustained performance over time is favored over rapid resource turnover.

Moreover, the study revealed significant interactive effects of transect and elevation on various traits, indicating that trait variation along elevational gradients is not uniform but varies among latitudinal transects. This complexity in plant responses to climatic gradients underscores the non-linear and multifaceted nature of plant adaptation. Overall, the results suggest that central populations tend to perform better than their marginal counterparts, potentially reflecting an environmental optimum. Rather than implying more efficient resource allocation, this optimum may result from moderate environmental conditions such as intermediate temperatures and adequate soil moisture that support greater resource availability and physiological functioning. Enhanced growth in these populations may therefore reflect favorable environmental inputs or a growth-maximizing strategy, rather than increased efficiency per se. These findings align with ecological theories that propose optimal performance at intermediate environmental gradients (Abeli et al., 2014), where conditions are neither too harsh nor too benign, allowing plants to optimize resource use and maintain resilience in the face of varying climatic conditions.

### Differential radial growth responses to recent climate change among populations

We also aimed to understand how *S. flabellaris* populations have responded to climate change in recent decades across various latitudinal transects and elevations and whether their climate sensitivity is consistent. Our results suggest that climate sensitivity varies with elevation but remains consistent across latitudinal transects. The pattern suggests that climatic changes, such as rising temperatures and altered precipitation patterns, drive growth increases, particularly at higher elevations where reduced low-temperature filtering allows for enhanced growth. This pattern also aligns with the globally observed phenomenon of elevation-dependent warming, where higher elevations are experiencing more rapid temperature increases (Pepin et al., 2015; Sharma et al., 2024). Such warming may extend the growing season and improve resource availability at upper elevation limits, thereby facilitating increased growth in these populations. Although there are no previous studies with a similar design, past research has reported population-specific climate sensitivity in plant populations (Bhuta et al., 2009; Fréjaville et al., 2020). We anticipated that lower elevations, which are warmer and face water limitation, would exhibit decreased growth due to increased water stress, as documented previously (Gazol et al., 2015). However, our results revealed no response in growth in low elevations likely resulting from the increased temperature being compensated for by increasing early summer precipitations, which have been reported from Western Himalayas(Sabin et al., 2020).

At higher elevations, low temperatures typically restrict plant growth (Körner, 2003). The observed growth increase at these elevations aligns well with expectations and can be attributed to a reduction in low-temperature filtering, allowing for greater growth. Mid-elevation populations showed no growth enhancement, possibly because these populations are already experiencing optimal growth conditions, leaving little room for further increase. Increased growth of dwarf shrubs like *S. flabellaris* could threaten other species in high elevation Himalayan ecosystems by altering competitive dynamics. The enhanced growth at higher elevations indicates that this species might occupy climatic niches previously unavailable, potentially disrupting the entire alpine ecosystem’s functioning. This shift could pose a risk to many species that are slower to respond to climate change. The differential growth responses among elevations also highlight the need for targeted management strategies to preserve biodiversity and ecosystem functioning in these sensitive regions.

## CONCLUSIONS

Our research identifies several key dimensions of trait variation in an alpine dwarf shrub, including hydraulic safety-conductivity trade-offs in wood anatomy, construction costs versus mechanical strength in twigs, and the exploitative-conservative strategy in leaves. These findings underscore the complex interplay of plant traits that contribute to overall performance and fitness. Our results indicate that central populations exhibit optimal performance. Climatic factors particularly growing season temperature and soil moisture, significantly influence stem anatomy, as well as the functional traits of twigs and leaves. Furthermore, recent climate warming has impacted populations differently across elevations, resulting in the most substantial growth increases at higher elevations. Significant intraspecific trait variations and population specific growth changes emphasize the need for site-specific differentiation when predicting the impacts of climate change on species performance and range shifts.

## AUTHOR CONTRIBUTIONS

Study design DT with help from ZM and JD: Field Data Collection: DT, NR; Laboratory Sample Processing and Analysis: DT, VJ, NR; Data Analysis: DT with help from JD; Writing – Original Draft Preparation: DT with editing and suggestions from JD; Writing – Review & Editing: DT, NR, JD, ZM; Funding Acquisition: DT, JD, ZM; Approval: All

## ACKNOWLEDGEMENTS

This work was funded by the European Union’s Horizon 2020 research and innovation programme under the Marie Sklodowska-Curie grant agreement No 101038052. The work was also partly supported by the Czech Science Foundation (projects no. 21-26883S and 24-11954S) and institutional funds of the Institute of Botany of the Czech Academy of Sciences (Grant Number: RVO 67985939).

## DATA AVAILABILITY

The data supporting this study’s findings will be submitted to an appropriate repository (Dryad/Zenodo).

## SUPPORTING INFORMATION

**Supplementary Table S1:** Characteristics of the localities studied along with climatic characteristics. MAT = mean annual temperature; MAP = mean annual temperature; GSSL = Growing season soil volumetric moisture content. Variables marked as DL are based on actual measurements using dataloggers. The growing season represents May to September. MAP and MAXT are based on data from CHELSA from 1999 to 2018.

| Latitudinal Transect | Elevation | MAP (mm) | MAXT (°C) | MAT<sup>DL</sup> (°C) | GST<sup>DL</sup> (°C) | GSSM<sup>DL</sup> (Volumetric) |
| --- | --- | --- | --- | --- | --- | --- |
| Northern | low | 462 | 3.29 | 3.83 | 9.53 | 0.233 |
|  | mid | 443 | 3.20 | 2.31 | 8.08 | 0.198 |
|  | high | 433 | 2.96 | 2.08 | 7.42 | 0.311 |
| Intermediate | low | 588 | 5.36 | 4.98 | 12.24 | 0.239 |
|  | mid | 575 | 5.01 | 3.54 | 10.56 | 0.238 |
|  | high | 565 | 4.84 | 2.22 | 7.66 | 0.301 |
| Southern | low | 1228 | 11.01 | 4.92 | 10.97 | 0.377 |
|  | mid | 1177 | 10.65 | 3.64 | 10.07 | 0.355 |
|  | high | 1172 | 10.55 | 3.04 | 9.64 | 0.363 |

**Supplementary Figure S1:**
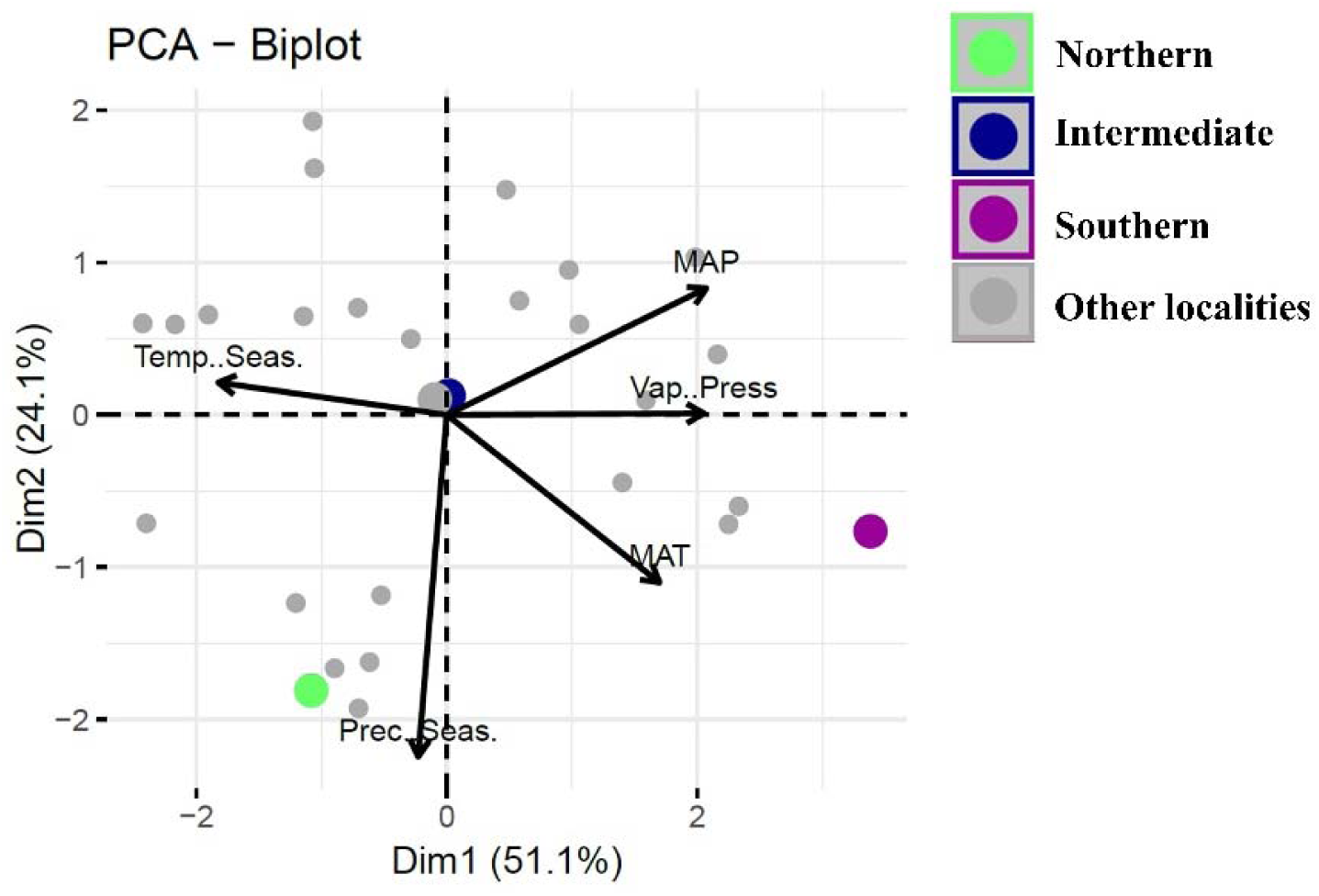
PCA plot showing the position of the latitudinal transects in the multivariate climatic space of *Salix flabellaris* among other known localities from Western Himalaya. MAT = Annual Mean Temperature, Temp.Seas. = Temperature Seasonality, MAP = Annual Precipitation, Prec.Seas. = Precipitation seasonality

**Supplementary Figure S2:**
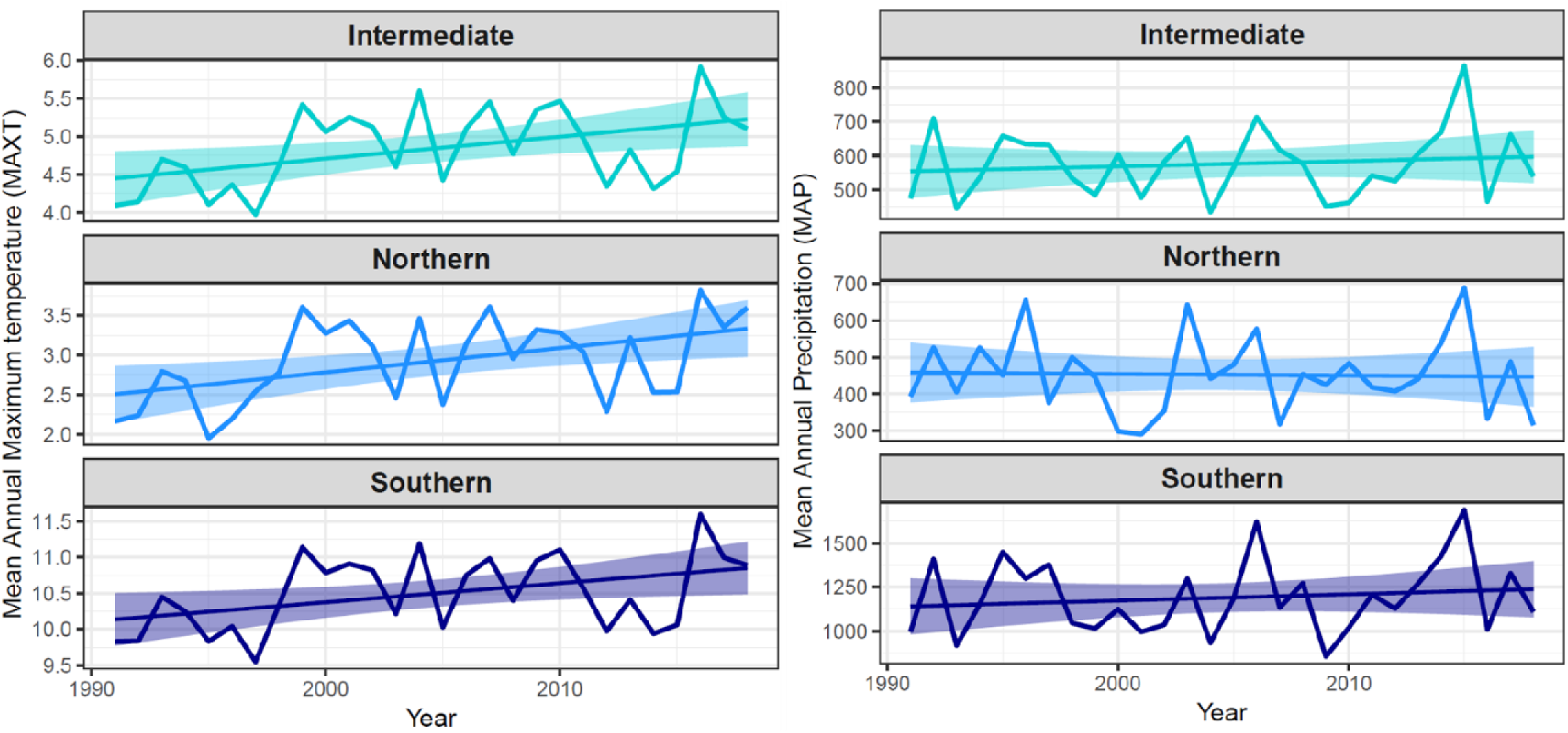
Temporal changes of mean annual maximum temperature (left) and mean annual precipitation (right) among different latitudinal transects in the recent past. Figures are based on data downloaded from CHELSA. Shaded areas represent 95% confidence intervals.

**Supplementary Figure S3:**
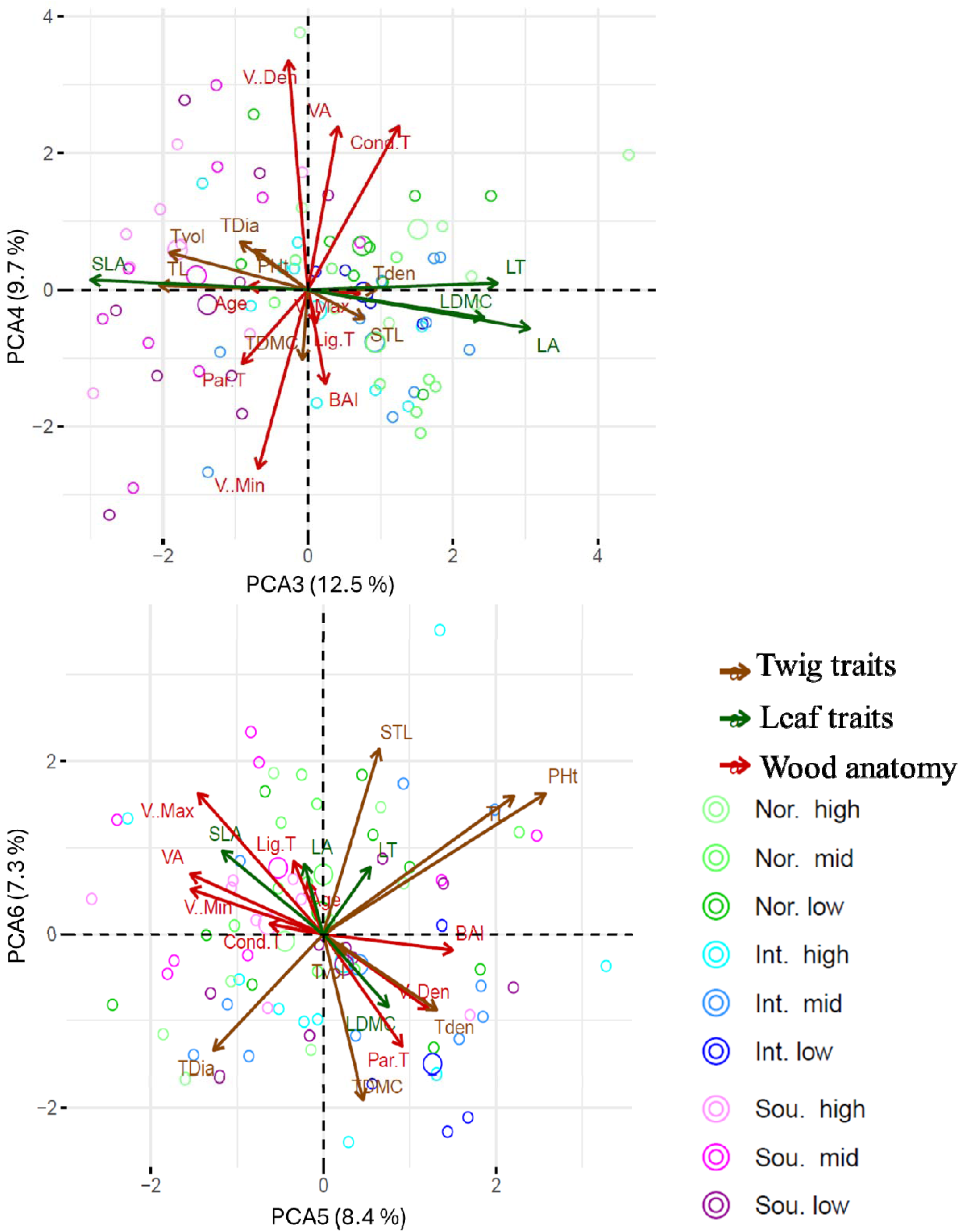
Principal component analysis (PCA) biplots showing the covariation of various traits in multivariate space. Axis 3 to 6 are presented in this figure and the axis 1 and 2 are presented in the main test in Figure 1. Int. = intermediate; Nor. = Northern; and Sou. = Southern transect. High, mid and low are the three elevations in each transect.

**Supplementary Figure S4:**
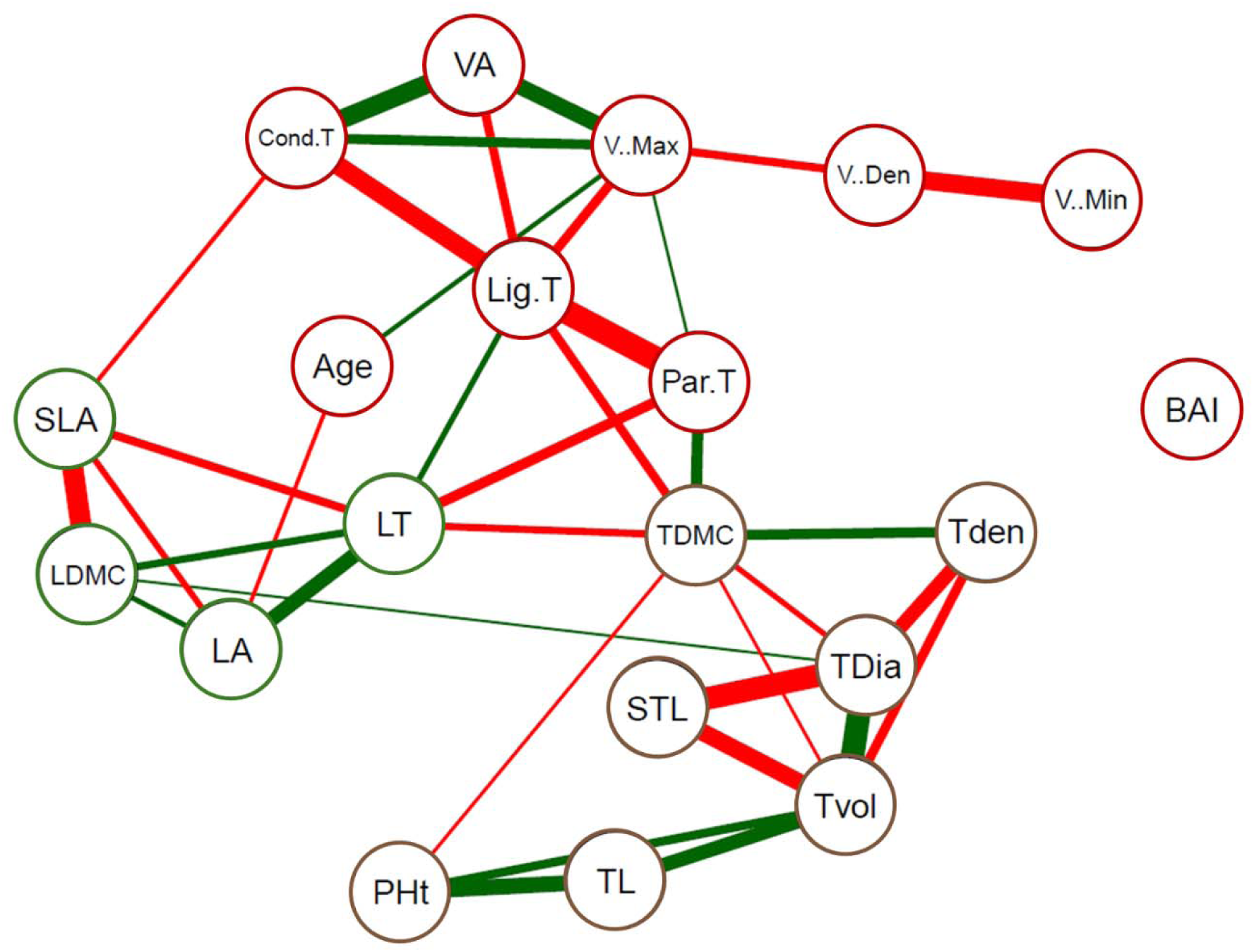
Correlations among different traits. Green colored connections among traits represent positive and red represents negative correlation. Thickness of line represents the strength of correlation. Non-significant correlations are not shown.

**Supplementary Figure S5:**
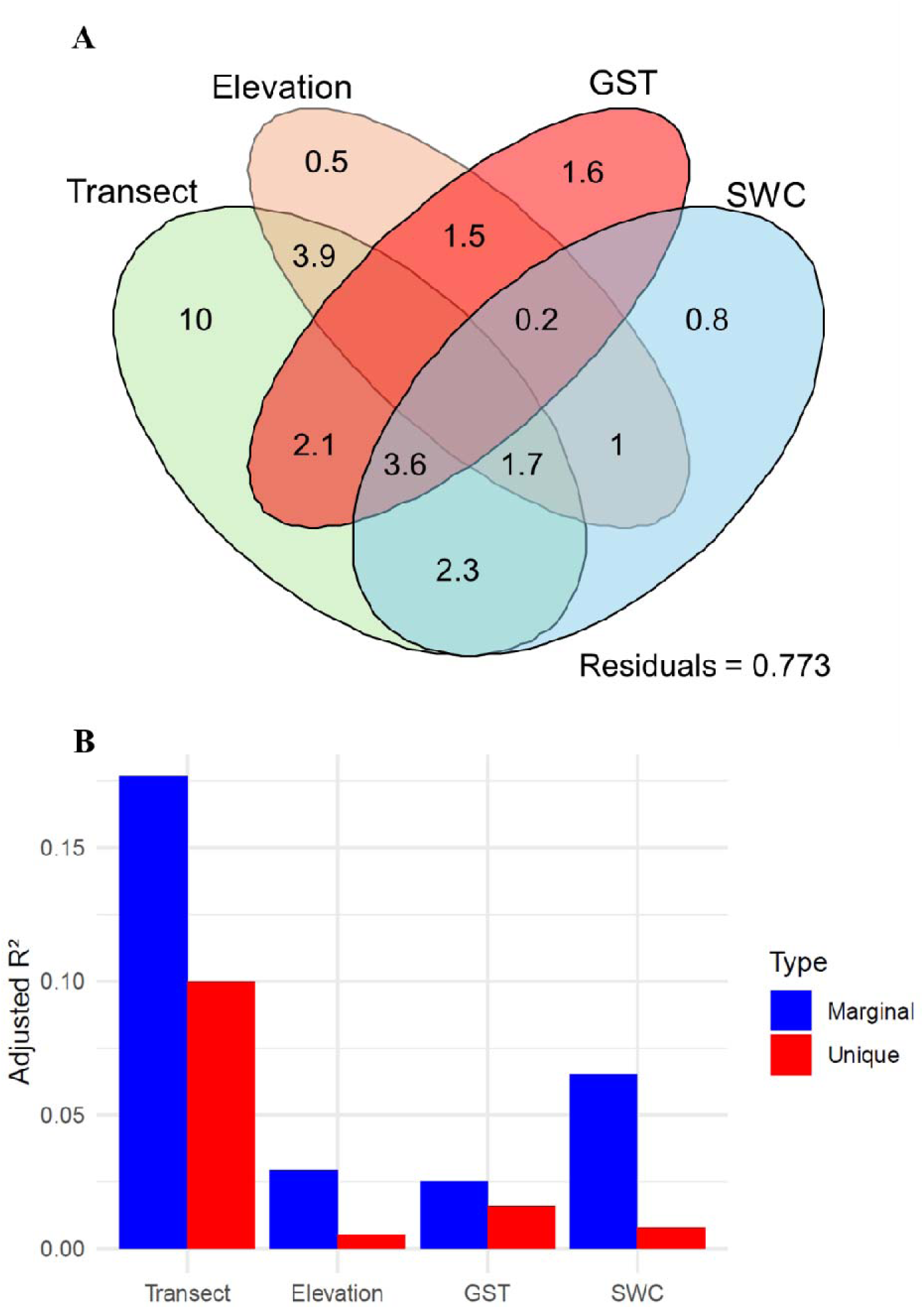
(A) Venn diagram illustrating the shared and unique variance in plant attributes explained by transect, elevation, growing season temperature (GST), and soil water content (SWC). (B) Bar plot showing the adjusted marginal and unique R² values for each predictor, quantifying their contributions to variation in plant attributes.

**Supplementary Figure S6:**
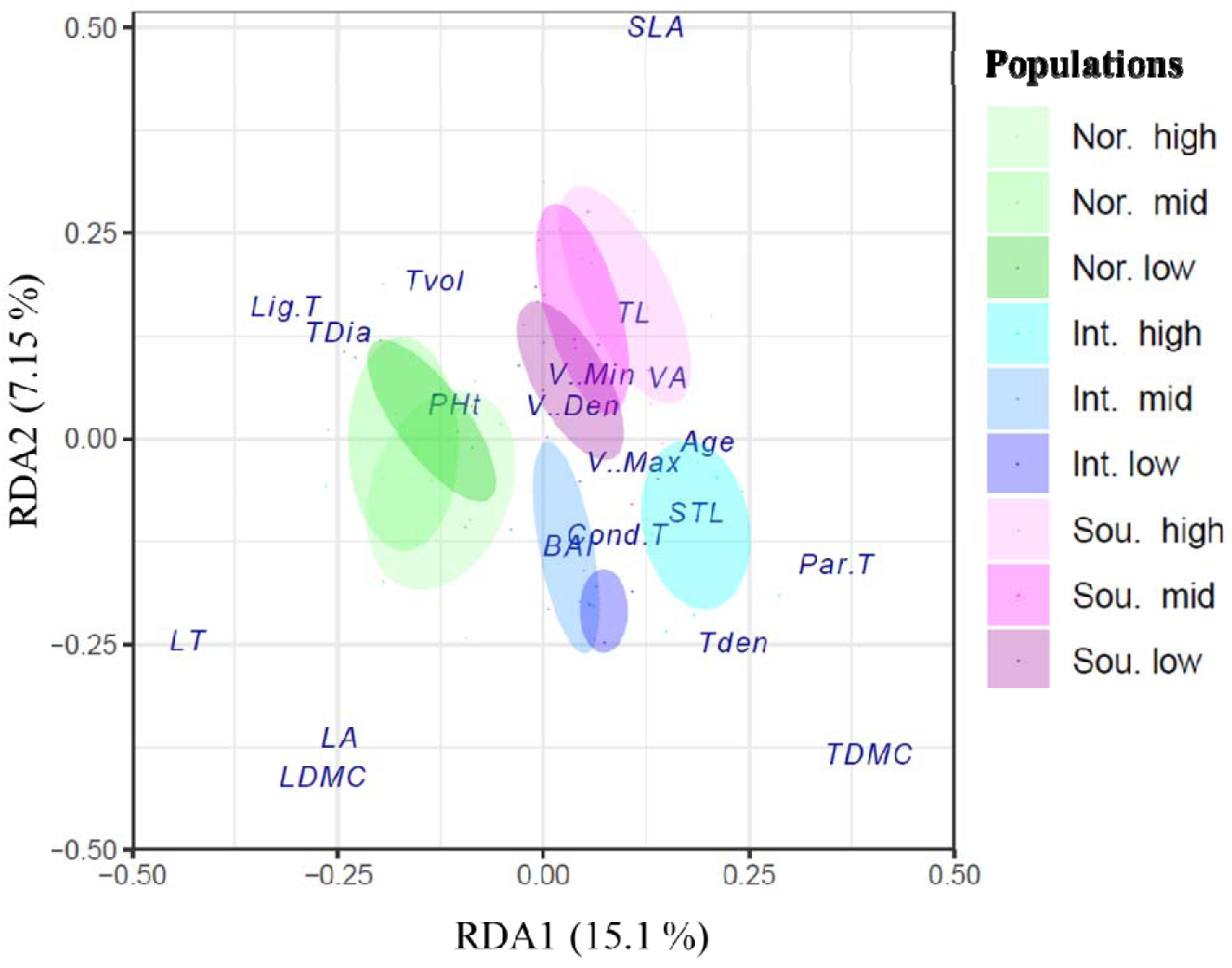
Redundancy analysis plots showing the effect of transect and elevation. Shaded areas represent 50% confidence intervals around the centroid.

